# Structurally conserved domains between flavivirus and alphavirus fusion glycoproteins contribute to replication in mammals and infectious virion production

**DOI:** 10.1101/2021.06.22.449399

**Authors:** Margarita V. Rangel, Nicholas Catanzaro, Sara A. Thannickal, Kelly A. Crotty, Maria G. Noval, Katherine E.E. Johnson, Elodie Ghedin, Helen M. Lazear, Kenneth A. Stapleford

## Abstract

Alphaviruses and flaviviruses have class II fusion glycoproteins that are essential for virion assembly and infectivity. Importantly, the tip of domain II is structurally conserved between the alphavirus and flavivirus fusion proteins, yet whether these structural similarities between virus families translate to functional similarities is unclear. Using *in vivo* evolution of Zika virus (ZIKV), we identified several novel emerging variants including an envelope glycoprotein variant in β-strand c (V114M) of domain II. We have previously shown that the analogous β-strand c and the ij loop, located in the tip of domain II of the alphavirus E1 glycoprotein, are important for infectivity. This led us to hypothesize that flavivirus E β-strand c also contributes to flavivirus infection. We generated this ZIKV glycoprotein variant and found that while it had little impact on infection in mosquitoes, it reduced replication in human cells and mice, and increased virus sensitivity to ammonium chloride, as seen for alphaviruses. In light of these results and given our alphavirus ij loop studies, we mutated a conserved alanine at the tip of the flavivirus ij loop to valine to test its effect on ZIKV infectivity. Interestingly, this mutation inhibited infectious virion production of ZIKV and yellow fever virus, but not West Nile virus. Together, these studies show that structurally analogous residues in the alphavirus and flavivirus class II fusion proteins contribute to virus infection *in vivo* and highlight these shared domains as targets for broad-spectrum arbovirus therapies.

**Author Summary:** Arboviruses are a significant global public health threat, yet there are no antivirals targeting these viruses. This problem is in part due to our lack of knowledge on the molecular mechanisms involved in the arbovirus life cycle. In particular, virus entry and assembly are essential processes in the virus life cycle and steps that can be targeted for the development of antiviral therapies. Therefore, understanding common, fundamental mechanisms used by different arboviruses for entry and assembly is essential. In this study, we show that flavivirus and alphavirus residues located in structurally conserved and analogous regions of the class II fusion proteins contribute to common mechanisms of entry, dissemination, and infectious virion production. These studies highlight how class II fusion proteins function and provide novel targets for development of antivirals.

## Introduction

Arthropod-borne viruses (arboviruses) are a diverse group of pathogens that can cause explosive epidemics and devasting disease [1–4]. Arboviruses can be transmitted by a wide variety of arthropod vectors including mosquitoes, ticks, and sandflies [5]. Importantly, vectors such as *Aedes* (*Ae.*) species of mosquitoes can transmit several pathogenic arboviruses such as dengue virus, Zika virus (ZIKV), and chikungunya virus (CHIKV), suggesting there may be common mechanisms of infectivity shared among specific arboviruses. Currently there are limited vaccines and no specific antiviral therapies targeting these viral threats, highlighting the need to better understand at the molecular level how these viruses replicate and are transmitted. In particular, identifying common mechanisms shared among arboviruses and genetically distant arbovirus families could help determine broad-spectrum characteristics needed for effective antivirals.

A common feature shared by all arboviruses is their need to be transmitted from an insect vector to the human host, yet we understand little of the molecular mechanisms involved in arbovirus transmission and infectivity. To study what contributes to arbovirus transmission we can look at West Nile virus (WNV) [6, 7], ZIKV [8], Venezuelan equine encephalitis virus [9, 10], and CHIKV [11]. Natural epidemics of these viruses have identified key residues in the viral glycoproteins that promote virus transmission and infectivity. These findings along with years of extensive experimentation have defined the viral glycoproteins as critical factors for virus assembly, attachment and entry, transmission, and pathogenesis [12–18].

Alphaviruses and flaviviruses encode class II fusion proteins that are required for pH-dependent entry and fusion [19, 20]. In a previous study, we used the *in vivo* evolution of CHIKV during mosquito-to-mammal transmission to identify and study key determinants for alphavirus infectivity and pathogenesis. In this study, we identified an adaptive variant (V80I) in β-strand c of the E1 glycoprotein domain II which contributes to CHIKV transmission in mosquitos and pathogenesis in mice [21]. In subsequent studies, we further characterized this residue and found that V80 functions together with residue 226 on the ij loop of the tip of domain II to impact pH- and cholesterol-dependent entry [22]. Importantly, the tip of domain II of the class II fusion glycoprotein is structurally conserved between alphaviruses and flaviviruses. This observation suggests that while these viruses are genetically unrelated, structural conservation in their glycoproteins could translate to functional similarities between virus families.

In this study, we used *in vivo* evolution of ZIKV to identify factors involved in flavivirus replication and infectivity. We identified novel variants in NS2A, NS3, and in β-strand c of domain II of the ZIKV envelope protein. Given the structural similarities between domain II of the alphavirus and flavivirus fusion glycoprotein, we hypothesized that β-strand c and the ij loop may have analogous functions in both virus families. Here, we found that a highly conserved valine in the ZIKV β-strand c attenuated virus replication in A549 cells and mice, and impacted pH-dependent entry. Moreover, we found that analogous residues in the tip of the envelope ij loop were important for ZIKV and yellow fever virus infectious virion production but not for WNV. Together, these data provide functional evidence that alphaviruses and flaviviruses use similar mechanisms and structural domains for infectivity and infectious virion production. These studies further our understanding of arbovirus biology as well as open new avenues for the development of antivirals targeting multiple virus families.

## Results

### Identification of emerging ZIKV variants in the envelope, NS2A, and NS3 during vector-borne transmission to type I interferon-deficient mice

In a previous study to understand how transmission route and organ microenvironment impacts ZIKV evolution, we dissected the *in vivo* evolution of the prototype ZIKV strain (MR766) in type I interferon-deficient mice (*Ifnar1^-/-^*) [23]. To do this, we infected *Ifnar1^-/-^* mice either by needle inoculation in the footpad or by mosquito bite from ZIKV infected *Ae. aegypti* mosquitoes, harvested organs at seven days post infection, and analyzed ZIKV variants by deep sequencing. Using the data from this study, we analyzed the synonymous and nonsynonymous minority variants from mice after mosquito transmission (**Fig. 1**). We identified unique variants in two mice (Mouse A and Mouse B) that had the same point mutations in multiple organs (**Fig. 1A and B**, **boxes**). In Mouse A, there were two synonymous changes (C1690T (8.2-12.9%) and T3931C (9.4-17.0%)). In Mouse B, there were three nonsynonymous changes (E: G1316A, V114M (8.9-9.2%); NS2A: A3638G, I33V (13.1-16.8%); and NS3: G4788A, R59K (19.4-22.7%). When we mapped these changes onto the protein structures we found that the E^V114M^ mutation is located in the tip of domain II which coincides with a highly conserved glycerophospholipid binding pocket [24] (**Fig. 1C**) and the R59K mutation in NS3 maps near a cellular 14-3-3 binding site [25] (**Fig. 1D**). Finally, the I33V mutation in NS2A is located in a putative transmembrane alpha-helix predicted to be in the ER lumen [14] (**Fig. 1E**).

**Fig. 1:**
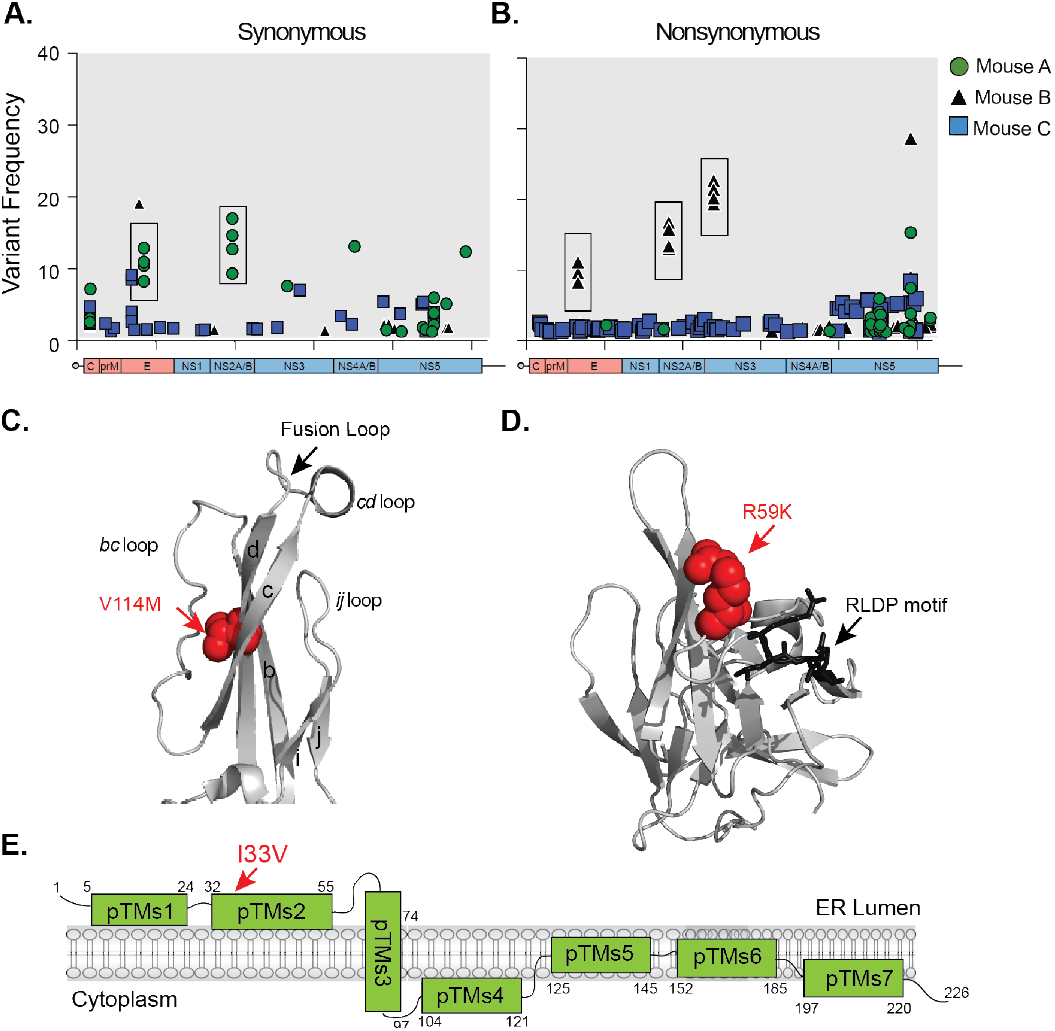
Identification of emerging ZIKV minority variants after vector-borne transmission. ZIKV MR766 vector-transmitted synonymous (**A**) and nonsynonymous) (**B**) minor variants (<50%) present in individual mice (N=3). Boxed variants signify variants present in multiple organs of individual mice. (**C**). Structure of the tip of ZIKV envelope protein (PDBID: 5JHM) with the V114M variant denoted in red. (**D**). Structure of the ZIKV NS3 protease (PDBID: 5GJ4) with R59K mutation in red and 14-3-3 RLDP binding motif in black. (**E**). Schematic representation of the ZIKV NS2A protein. The I33V mutation is depicted in red.

### ZIKV E, NS3, and NS2A variants have no significant advantage in *Ifnar1^-/-^* mice and are attenuated in A549 cells

Given that these ZIKV variants were identified in *Ifnar1^-/-^* mice, we tested whether this variant had a replication advantage in *Ifnar1^-/-^* mice. We introduced each variant individually into an infectious clone of ZIKV strain MR766, infected *Ifnar1^-/-^* mice with each virus, and monitored survival and weight loss. All four viruses caused 100% lethality, with no significant difference in the average time to death (**Fig. 2A**). Interestingly, we found that although mice infected with each variant succumbed to infection at the same time as wild-type (WT) ZIKV, the mice infected with the NS2A^I33V^ and NS3^R59K^ variants initially gained weight followed by rapid weight loss and death (**Fig. 2B, green and purple triangles**). These data suggest that NS2A and NS3 may play critical roles in ZIKV pathogenesis later in infection.

**Fig. 2:**
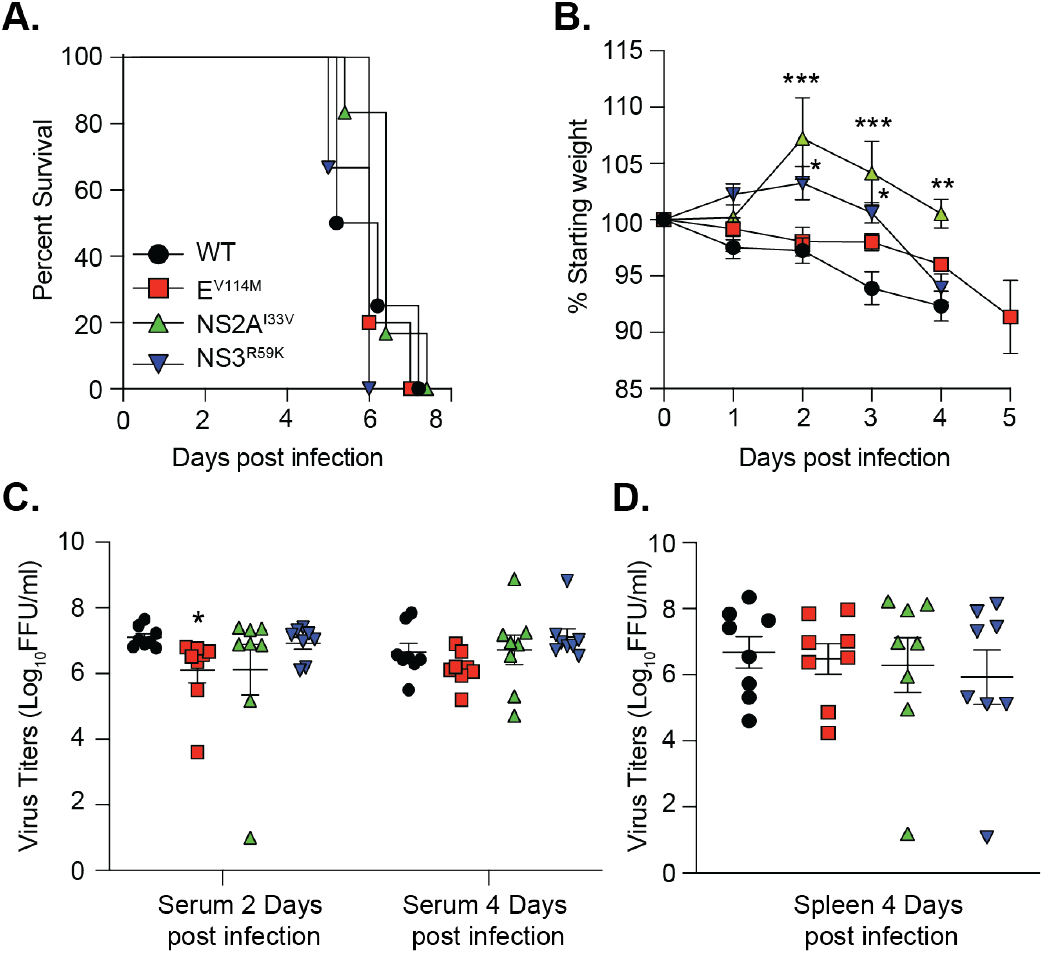
ZIKV variants impact weight loss and virus replication in *Ifnar1^-/-^* mice. (**A**). Survival of 5- to 6-week-old *Ifnar1^-/-^* mice infected with 10^3^ FFU of wild-type (WT) (N=4), E^V114M^ (N=5), NS2A^I33V^ (N=6), or NS3^R59K^ (N=6) ZIKV. No significant difference between groups (**B**). Weight loss of mice in (**A**). Data are censored after the first mouse in each group died. Two-way ANOVA. *** p<0.001, * p<0.05. Viral loads in serum (**C**) and spleen (**D**) from *Ifnar1^-/-^* mice infected with WT ZIKV or variants. N=8 for each virus. One-way ANOVA, Kruskal-Wallis test. * p<0.05.

To investigate differences in viral replication between variants, we infected *Ifnar1^-/-^* mice with each variant and quantified infectious virus in the serum after 2- and 4-days post-infection, and in the spleen at 4-days post infection (**Fig. 2C and D**). We found no significant advantage of any of these variants in *Ifnar1^-/-^* mice and, if anything, there was a reduction in replication consistent with the weight loss results. Finally, to investigate the replication of each ZIKV variant in cell culture, we performed multi-step growth curves in human A549, monkey Vero, and *Ae. albopictus* C6/36 cells. We found that ZIKV E^V114M^ and NS2A^I33V^ were significantly attenuated in A549 cells compared to WT virus (**Fig. 3A**), yet showed no differences in replication in Vero or C6/36 cells (**Fig. 3C and D**). Together, these data indicate that these ZIKV variants show no significant replication advantage in *Ifnar1^-/-^* mice, yet are attenuated in IFN-competent human cells, highlighting a role for these residues in ZIKV replication.

**Fig. 3:**
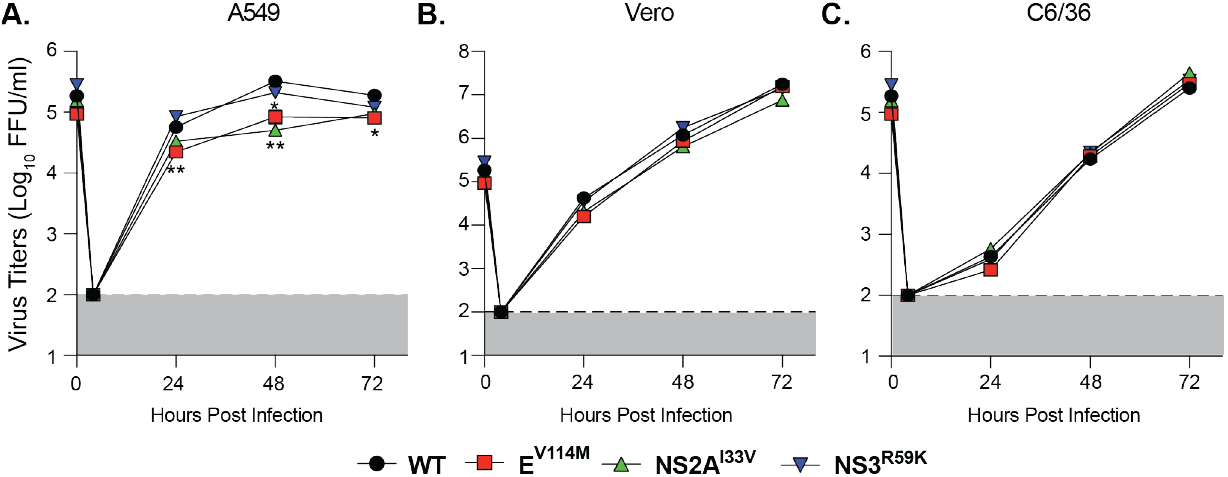
Replication of ZIKV envelope and NS2A variants is attenuated in A549 cells. Multi-step growth curves of WT ZIKV and variants in A549 (**A**), Vero (**B**), and C6/36 cells (**C**). Each cell line was infected with each virus at an MOI = 0.1 and virus titers in the supernatant were quantified by focus-forming assay. Dotted lines indicates the limit of detection. N=3 independent experiments. Two-way ANOVA. **p<0.01.

### ZIKV E residue V114 is structurally analogous to alphavirus E1 residue V80 and is important for pH-dependent entry

We previously identified the CHIKV E1 glycoprotein residue V80 in β-strand c as important for CHIKV pH-dependent entry [22]. Interestingly, the ZIKV envelope residue V114 falls into a structurally analogous site in β-strand c of the ZIKV E protein (**Fig. 4A**), and similar to alphaviruses [22], a valine at flavivirus residue 114 is highly conserved (**Fig. 4B**). Given the similarities between ZIKV E^V114^ and CHIKV E1^V80^, we hypothesized that ZIKV E^V114^ may also contribute to ZIKV pH-dependent entry. To test this hypothesis, we first tested that the ZIKV E^V114M^ variant did not lead to any major differences in viral protein accumulation or processing. To address this, we transfected 293T cells with infectious clones encoding WT ZIKV, a replication-dead ZIKV harboring a mutation (GNN) in the polymerase, or the E^V114M^ variant. We harvested cells at 72 hours post-transfection and evaluated the accumulation of ZIKV proteins by western blot. While we found a slight reduction in all proteins in the E^V114M^ variant, possibly due to the replication defect in human cells, there were no major differences in protein processing with the E^V114M^ variant (**Fig. 4C**). Next, we asked whether the ZIKV E^V114M^ variant was sensitive to the lysosomotropic agent ammonium chloride which will deacidify the endosome and block pH-dependent entry. We infected Vero cells with each virus in the presence of increasing concentrations of ammonium chloride and found that ZIKV E^V114M^ replication was significantly more sensitive to the presence of ammonium chloride, mimicking what we observed with CHIKV E1^V80^ variants [22] (**Fig. 4D**). Together, these data indicate that the flavivirus glycoprotein β-strand c contributes to pH-dependent entry as seen for alphaviruses.

**Fig. 4:**
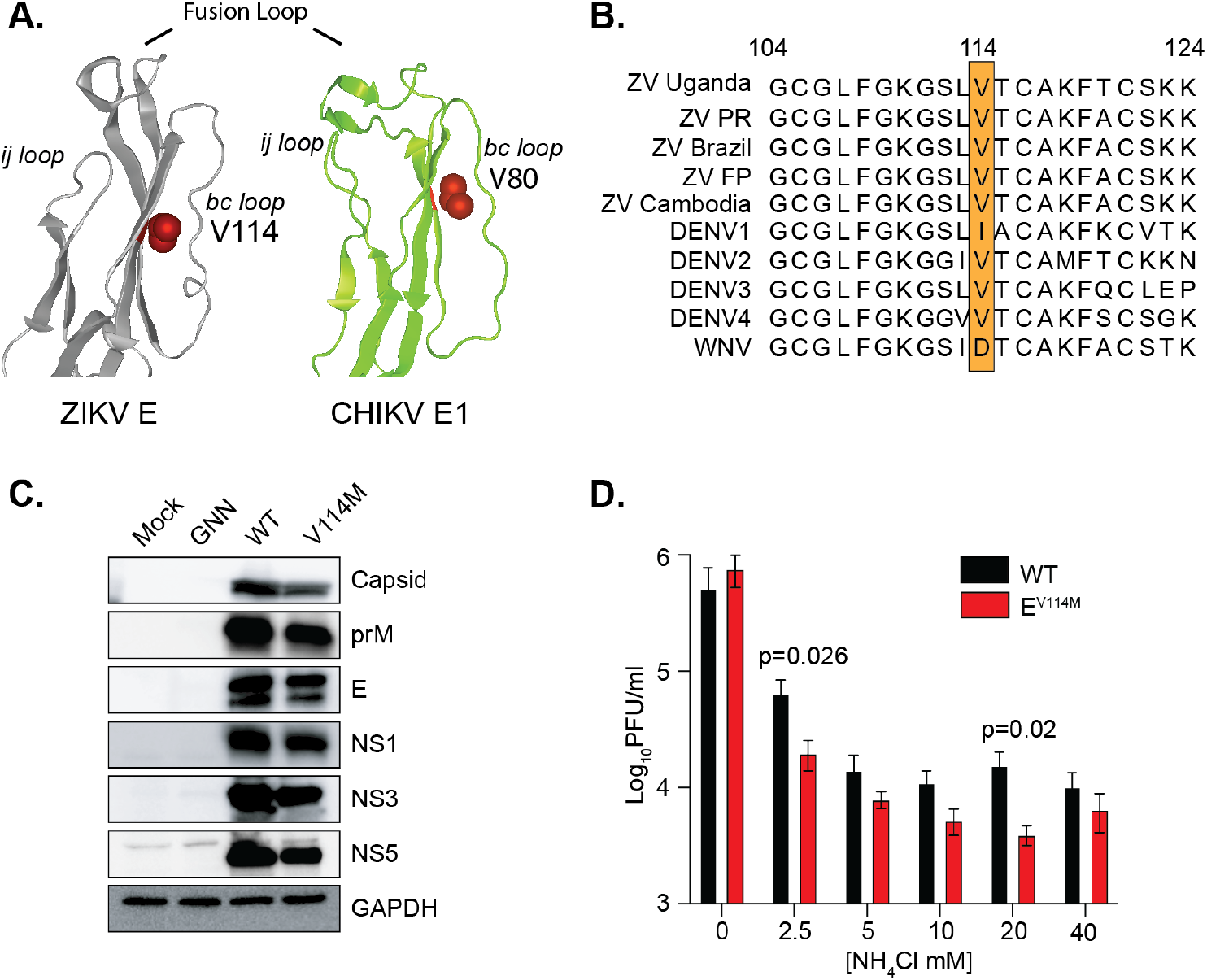
ZIKV V114M is structurally analogous to chikungunya virus E1 V80 and sensitive to ammonium chloride inhibition. (**A**). ZIKV MR766 envelope (PDBID: 5JHM) and CHIKV E1 (PDBID: 3n42). The fusion loop, ij loop, bc loop, and V114 and V80 variants are denoted in red. (**B**). Flavivirus envelope protein sequence alignment around residue V114. (**C**). 293T cells were transfected with a ZIKV WT or ZIKV E^V114M^ infectious clone. Cells were harvested at 72 hpi, lysed in Laemmli buffer, and ZIKV proteins were analyzed by SDS-PAGE and immunoblotting. Blot is a representative of at least three independent transfections. (**D**). Vero cells were pretreated for 1 hr with increasing concentrations of NH4Cl and infected at MOI = 0.1. Supernatant was collected 36 hpi and viral titers were quantified by plaque assay. Data represent the mean and SEM. N=3 independent experiments with internal technical triplicates. Students *t*-test.

### ZIKV E^V114M^ is attenuated in wild-type neonatal mice with no impact in replication in *Ae. aegypti* mosquitoes

We observed that CHIKV E1^V80^ contributes to dissemination in wild-type mice and in *Ae. aegypti* mosquitoes [22]. Therefore, we hypothesized that ZIKV E^V114M^ and ZIKV envelope β-strand c may also play essential roles in dissemination in mice and mosquitoes. To study ZIKV replication in WT immunocompetent mice, we first infected 6-week-old C57BL/6 mice; and as expected, neither virus replicated in these mice (data not shown). As an alternative approach, we used 4- and 7-day old C57BL/6 neonatal mouse models. We hypothesized that the 4-day old mice, which are more susceptible to ZIKV, would allow us to mimic the *Ifnar1^-/-^* mice while the 7-day old mice would allow us to study ZIKV infection in a model that is closer to adult mice. We infected each neonatal model with WT ZIKV or the ZIKV E^V114M^ variant. We found that ZIKV E^V114M^ was attenuated in its ability to replicate and disseminate in both 4- and 7-day old mice (**Fig. 5A and B**). This attenuation of ZIKV E^V114M^ was dependent on the age of the mice as infection of the ZIKV E^V114M^ variant was not detectable in the brain of several 7-day old mice, suggesting that the virus may be cleared faster than WT ZIKV or there is a subset of 7-day old mice that are specifically not susceptible to infection with the ZIKV E^V114M^ variant.

**Fig. 5:**
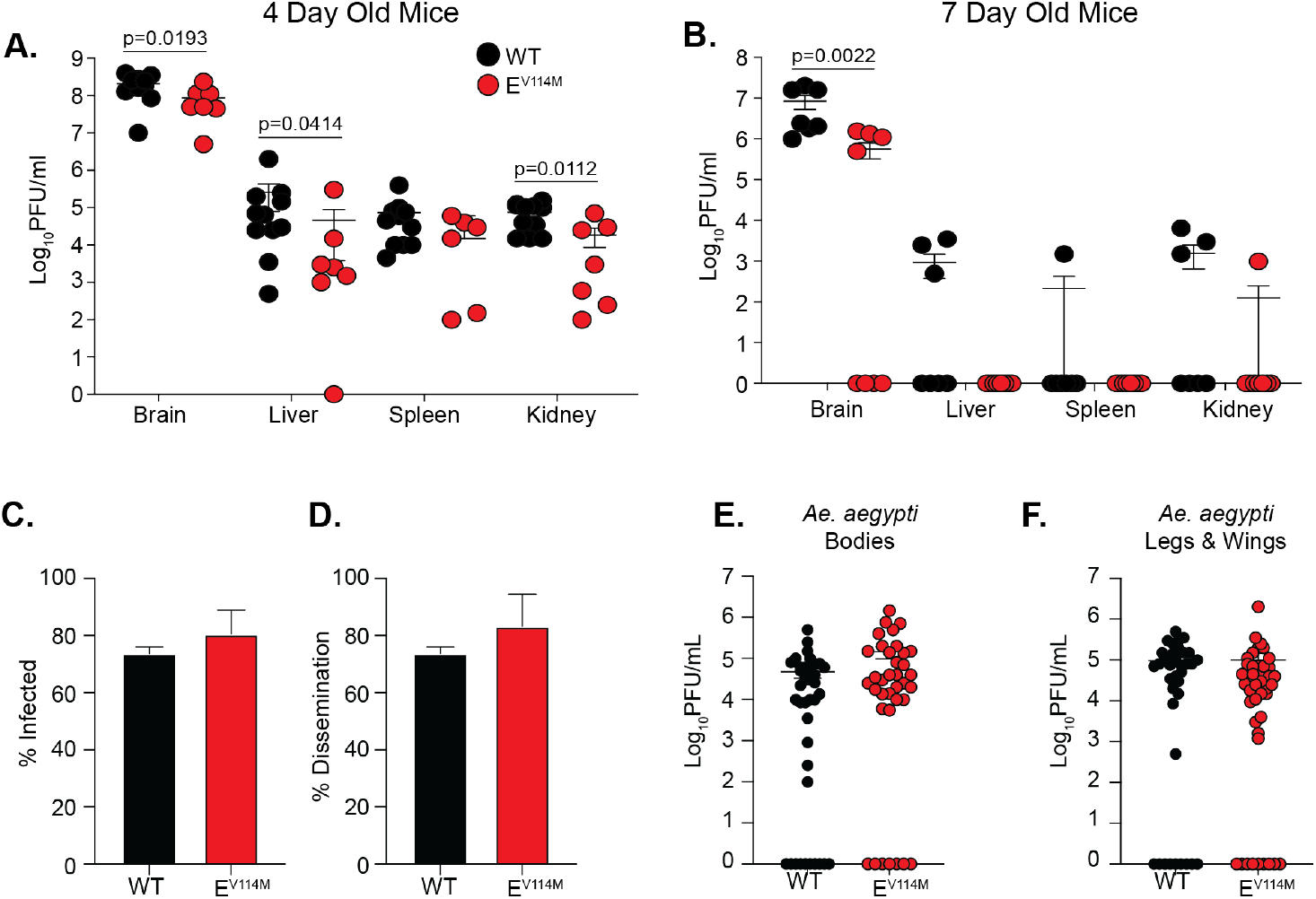
Replication of WT and E V114M ZIKV in neonatal mice and *Ae. aegypti* mosquitoes. 4 (**A.**) and 7 (**B.**) day old C57BL/6J mice were infected subcutaneously with 10^4^ PFU of each virus. Virus titers were quantified in each organ at 7 days post infection. Data represent the mean and SEM of two independent infections. 4 day old mice (WT N=11, E^V114M^; N=7). 7 day old mice (WT; N=7, E^V114M^; N=8). Mann-Whitney test. **C-F.** *Ae. aegypti* mosquitoes were infected with 10^6^ PFU of each virus and viral titers were determined in the bodies (**C. and E.**) and legs and wings (**D. and F.**) at 14 days post infection. Data represent the mean and SEM of two independent infections. WT; N=39, E^V114M^; N=36. No significant different between groups. Mann-Whitney compare ranks test.

To investigate how ZIKV E^V114M^ impacts replication in mosquitoes, we infected *Ae. aegypti* mosquitoes with WT ZIKV or the ZIKV E^V114M^ variant and quantified viral titers in the bodies, and legs and wings at 14 days post infection (**Fig. 5C-F**). We found no major difference in the percentage of infected mosquitoes (**Fig. 5C**) or the percentage of mosquitoes that had virus in the legs and wings (disseminated virus) (**Fig. 5D**). Interestingly, while we did not find a statistically significant increase in viral titers with the ZIKV E^V114M^ variant, the titers were higher in multiple mosquitoes with no clear replication disadvantage (**Fig. E and F**). Together, these data suggest that the flavivirus envelope β-strand c contributes to replication in neonatal mice with no negative impact on replication in mosquitoes.

### A conserved alanine at the tip of the flavivirus envelope protein ij loop is important for ZIKV infectious particle production

CHIKV has undergone several natural adaptation events that have shaped its infectivity and transmission [26–28]. One of the most significant was the emergence of a single amino acid substitution in the E1 glycoprotein (A226V) during an epidemic on La Reunion Island [11]. This variant and residue 226, located in the ij loop of the alphavirus E1 glycoprotein, has been shown to increase replication and transmission in *Ae. albopictus* mosquitoes [27, 29] and is important for cholesterol-dependent entry [30,31]. Interestingly, flaviviruses contain a highly conserved alanine at the tip of the ij loop similar to CHIKV (**Fig. 6A**). Given the structural similarities between the flavivirus and alphavirus class II fusion proteins, we hypothesized that the ZIKV alanine would be important for virus infectivity. We mutated this alanine (A249) to a valine in both the MR766 and Brazilian strains of ZIKV and found that this single mutation completely blocked the production of infectious ZIKV particles in 293T cells as well as in BHK-21 and Vero cells (**Fig. 6B,** data not shown). One potential explanation for this phenotype is that this mutation inhibits RNA replication and subsequent protein production, as is the case for the NS5 GNN mutant (**Fig. 6**). To address whether ZIKV E^A249V^ impacted replication and/or protein production, we transfected 293T cells with each ZIKV plasmids, harvested cells at 72 hours post-transfection, and analyzed protein accumulation by western blotting. We detected the majority of ZIKV proteins in transfected cells, albeit at lower levels due to the lack of spread of the ZIKV E^A249V^ variant (**Fig. 6C**). Interestingly, the envelope protein did not accumulate during transfection, suggesting that the virus is replicating yet may not be producing infectious particles due to a defect in envelope accumulation (**Fig. 6C**). These results indicate that the ZIKV ij loop is critical for infectious particle production potentially through the stability of the envelope glycoprotein.

**Fig. 6:**
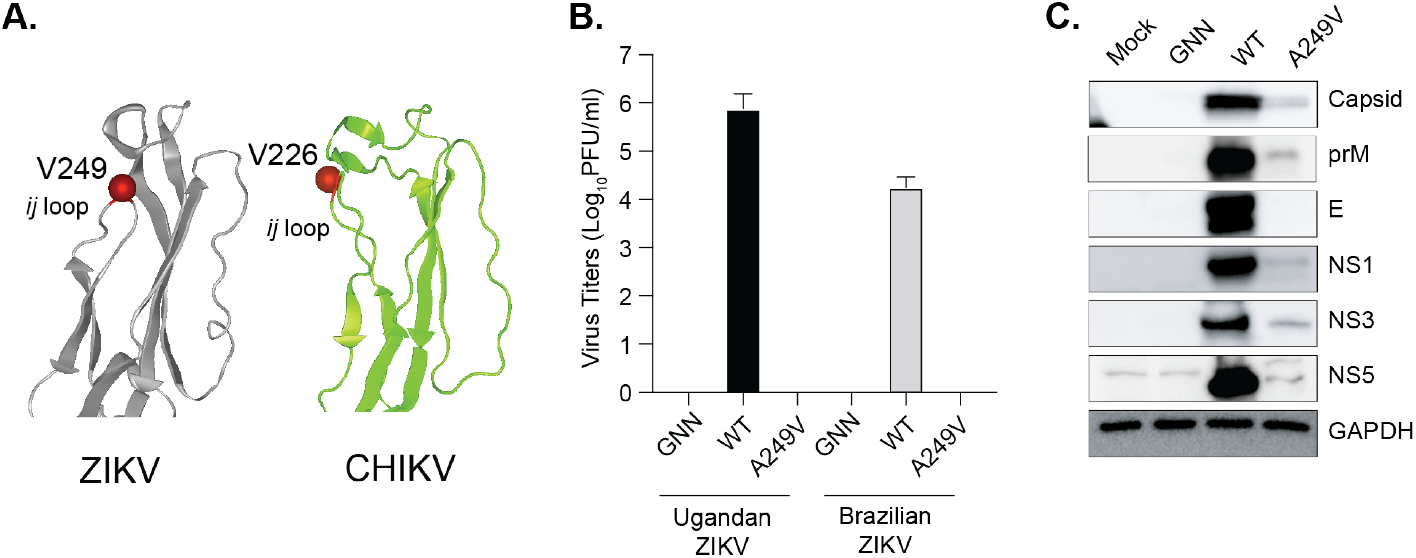
A single point mutant in the flavivirus E ij loop inhibits ZIKV infectious particle production and impacts envelope protein accumulation. (**A**) ZIKV MR766 envelope (PDBID: 5JHM) and CHIKV E1 (PDBID: 3n42). The ij loop and ZIKV A249 and CHIKV A226 residues are denoted in red. (**B**) 293T cells were transfected with WT ZIKV or ZIKV E^A249V^ infectious clone plasmids and virus supernatants were collected at 48 hpi. Virus titers were quantified by plaque assay. N=2 with internal technical duplicates. (**C**) 293T cells were transfected with WT ZIKV or ZIKV E^A249V^ plasmids. Cells were harvested at 72 hpi, lysed in Laemmli buffer, and ZIKV proteins were analyzed by SDS-PAGE and immunoblotting. Immunoblot is a representative of at least 3 independent transfections.

### Conserved alanine at the tip of flavivirus ij loop blocks YFV but not WNV infectious particle production

The alanine at the tip of the flavivirus ij loop is highly conserved among flaviviruses and sits in the middle of a histidine and lysine which have been shown to be important for virion assembly [32] (**Fig. 7A and B**). Given this conservation, we hypothesized that the ij loop could also be critical for infectious particle production of other flaviviruses. To test this hypothesis, we mutated the alanine at the tip of the E protein ij loop to a valine in the yellow fever virus (YFV) and WNV infectious clones, transfected Vero cells with *in vitro* transcribed RNA, and quantified infectious virions in the supernatant 48 hours later by plaque assay. We found that the A-to-V mutation in YFV significantly impaired infectious virus production, while this same mutation in WNV has no effect (**Fig. 7C and D**). Importantly, when we sequenced the infectious virus from WNV, the A247V mutation was maintained and genetically stable. These results indicate that the ij loop plays flavivirus-specific roles in virion assembly and the single alanine at the tip of the ij loop contributes to infectious particle production. Future work will be essential to understand how flavivirus and alphaviruses coordinate infections in insects and mammals and to elucidate the differences in virus entry between flaviviruses.

**Fig. 7:**
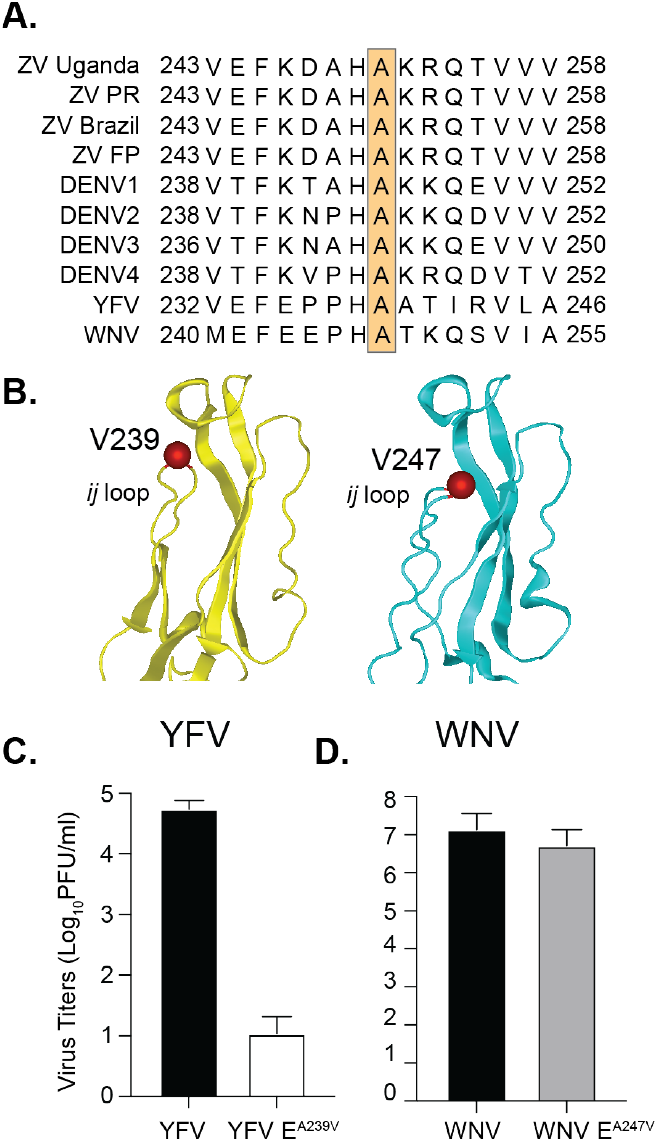
The alanine at the tip of flavivirus envelope inhibits infectious particle production of yellow fever virus but not West Nile virus. (**A**) Flavivirus envelope ij loop protein sequence alignment. Orange box indicates the conserved alanine at the tip of the ij loop. (**B**) Domain II of the YFV (PDBID: 6iw4) and WNV (PDBID: 2HG0) envelope protein. The ij loop and YFV A239 and WNV A247 residues are denoted in red. (**C**) Vero cells were transfected with WT or variant YFV and WNV *in vitro* transcribed RNA and virus containing supernatant was collected at 48 hpi. Virus titers were quantified by plaque assay. N=3 independent experiments.

## Discussion

Arboviruses are a diverse group of human pathogens belonging to multiple virus families. There are no antiviral therapies targeting these viruses. This problem is in part due to our incomplete understanding of the common, fundamental molecular mechanisms different arboviruses use for infectivity and spread. To identify and study the fundamental mechanisms of arbovirus biology, we use samples collected from natural infections and *in vivo* lab experiments. In this study, we observed the emergence of ZIKV variants during vector-borne transmission where three variants of interest were found in the E (V114M, ~10%), NS2A (I33V, ~15%), and NS3 (R59K, ~20%) proteins. The envelope variant is located at the tip of the class II fusion protein, near the fusion loop and the highly conserved glycerophospholipid-binding pocket [24]. The NS2A protein variant is located in a predicted alpha-helix near the ER membrane, and the NS3 protease variant is close to an RLPD motif that was recently shown to be important for interactions between NS3 and cellular 14-3-3 proteins to antagonize the innate immune response [25]. Given that we originally found these variants in *Ifnar1^-/-^* mice, we hypothesized that they would play significant roles in ZIKV infection in mice. However, when in separate experiments we introduced these variants into *Ifnar1^-/-^* mice, they had no significant replication advantage in the organs we investigated. These data and the fact that these variants were not found in mice infected via needle inoculation [23], suggests that these variants may not have been selected for in mice but rather in *Ae. aegypti* mosquitoes. This observation is particularly interesting as the alphavirus glycoproteins have been highlighted many times in nature and in the lab as a key evolutionary determinant for alphavirus infectivity and transmission in mosquitoes [21, 26, 28]. This study provides evidence that this may also be the case for flaviviruses as well. Importantly, future studies regarding the role of NS2A and NS3 in mosquitoes and mice will be important to better understand the molecular mechanisms of the flavivirus life cycle *in vivo.*

The ZIKV envelope has been shown to significantly contribute to virulence [8, 33–38]. The ZIKV envelope residue V114 is located in β-strand c of the glycoprotein domain II. Because β-strand c of CHIKV plays an important role in virus dissemination and pathogenesis in mice and transmission in mosquitoes, we hypothesize this would also be the case for flaviviruses. Interestingly, we found that the ZIKV E^V114M^ variant was attenuated in multiple mouse models and in A549 cells, confirming that E residue V114 and β-strand c is important for flavivirus infection. Given this attenuation in mammals, we thought that the ZIKV E^V114M^ would have a significant replication advantage in mosquitoes, yet while we did see an increased trend in replication of the ZIKV E^V114M^ variant in *Ae. aegypti* mosquitoes, this was not statistically significant. One potential explanation for this could be that the E^V114M^ mutation alone is not enough to enhance replication and perhaps the combination of E, NS2A, and NS3 variants will drive enhanced replication in mosquitoes and/or mice. Indeed, mutations in ZIKV NS3 have been implicated in ZIKV infection in mice [39]. Moreover, as we only looked at one late time point, it will be interesting to explore the temporal replication of these variants in mosquitoes and mice to understand where the variants are acting during the viral life cycle. This is particularly important for the NS2A and NS3 variants in mice, as we found that mice infected with these variants gain weight in the beginning of the infection and rapidly succumb to the infection, suggesting that NS2A and NS3 may play a role late in the infection.

In addition to the envelope domain II β-strand c, we also observed structural similarities between the alphavirus and flavivirus glycoprotein ij loop. In CHIKV, a mutation from alanine to valine at position 226, in the tip of the ij loop, was responsible for a significant outbreak that led to increased replication in *Ae. albopictus* mosquitoes and to the emergence of the Indian Ocean Lineage of CHIKV [11, 27]. However, when we made this same A-to-V mutation in ZIKV, there was complete inhibition of infectious particle production, suggesting that the ij loop may play a role in ZIKV particle assembly. This is plausible as previous work with Japanese encephalitis virus (JEV) has shown that mutation of a conserved histidine and lysine in the ij loop impacts JEV assembly [32]. In that study, the authors had hypothesized that these conserved charged His and Lys residues in the envelope ij loop interact with negatively charged residues in prM to promote assembly. This idea is intriguing. It is possible that the valine mutation in the ij loop also destabilizes interactions between E and prM leading to reduction in infectious particle production. However, this does not directly explain the specific reduction in the accumulation of the E^A249V^ protein in cells. Future studies will be important to understand how the ij loop contributes to E stability and how the ij loop impacts ZIKV assembly.

Finally, we found that the alanine at the tip of the flavivirus ij loop behaves in a flavivirus specific manner in that a mutation of alanine to valine in ZIKV and YFV blocked infectious virus production, yet there was no such effect with WNV. In comparing the crystal structures of the ZIKV, YFV, and WNV ij loop (**Fig. 7**) we noted that the alanine is oriented in a different direction in WNV; and perhaps this change in structure is responsible for this phenotype. On the other hand, there could be many other genetic differences between the envelope of ZIKV and WNV that account for these differences in infectious particle production. Importantly, residue 114 in ZIKV is a valine and in WNV 114 encodes for an aspartic acid, already suggesting differences in E β-strand c function. This may help shed light on how flaviviruses such as ZIKV and WNV infect different mosquito species and hosts such as birds.

Together, these studies have identified novel flavivirus variants that impact flavivirus infection. We have shown that structurally similar domains of the flavivirus and alphavirus envelope glycoproteins contribute to similar functions in virus infection. These common mechanisms of infection could potentially provide the determinants needed for the development of broad-spectrum antivirals. More importantly, these studies lead to new insight into the fundamental biology of flaviviruses. Future studies to dissect the molecular details of the flavivirus and alphavirus envelope proteins are critical to understand how these viruses function and cause disease.

## Materials and Methods

### Cells lines

293T cells (ATCC CRL3216) and Human Foreskin Fibroblasts (HFF1, ATCC SCRC-1041) were grown in Dulbecco’s Modified Eagle Medium (DMEM) supplemented with 1% penicillin/streptomycin (P/S), 1% nonessential amino acids, and 10% heat-inactivated fetal bovine serum (FBS, Atlanta Biologicals) at 37°C with 5% CO_2_. Stapleford Lab Vero cells (ATCC CCL-81) were maintained in DMEM supplemented with 1% P/S and 10% heat-inactivated newborn calf serum (Gibco) at 37°C with 5% CO_2_. Lazear Lab Vero and A549 cells were maintained in DMEM containing 5% heat-inactivated FBS and L-glutamine at 37°C with 5% CO_2_. C6/36 cells were maintained in DMEM containing 6% FBS, NEAA, and P/S at 28°C with 5% CO_2_. All cells were verified to be mycoplasma free.

### Viruses

The ZIKV (Ugandan 1947 – MR766) wild-type and replication-deficient (NS5 GNN mutant) plasmid infectious clones were a gift from Dr. Matthew Evans at the Icahn School of Medicine at Mt. Sinai [40]. The ZIKV Brazilian strain infectious clone was a gift from Dr. Alexander Pletnev at the National Institutes of Health [41]. The yellow fever virus (vaccine strain 17-D) infectious clone was a gift from Dr. Julie Pfeiffer at the University of Texas Southwestern [42]. The West Nile virus infectious clone was a gift from Dr. Gregory Ebel at Colorado State University [43]. The ZIKV, WNV, and YFV envelope variants were generated by site-directed mutagenesis using Phusion high-fidelity polymerase (Thermo) and the primers in **Table 1**. All plasmids were sequenced in full to confirm no second-site mutations were introduced.

**Table 1:**
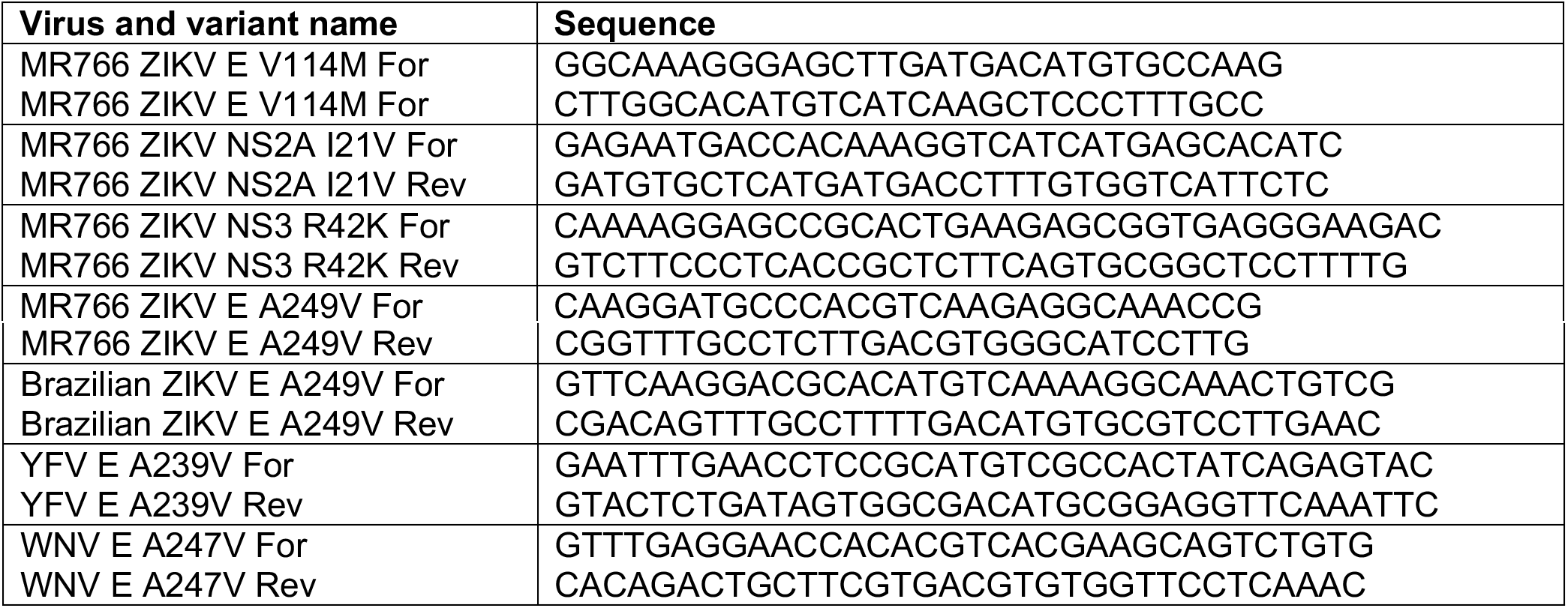
Primers used for this study.

The wild-type and variant Ugandan and Brazilian ZIKV were produced by transfecting 293T cells with 0.5 μg of each plasmid via Lipofectamine 2000 transfection reagent (Invitrogen). Virus-containing supernatants were harvested 48 hrs post transfection, centrifuged at 1,200 x g for 5 mins, aliquoted and stored at −80°C. To generate a working virus stock, Vero cells were infected with transfection virus stocks and virus-containing supernatants were harvested 48 hrs post infection, centrifuged at 1,200 x g for 5 mins, aliquoted, and stored at −80°C. Viral titers were determined by plaque assay on Vero cells, as described below.

Wild-type and variant YFV and WNV were generated by the transfection of *in vitro* transcribed RNA into Vero cells. In brief, YFV and WNV infectious clones were linearized overnight with XhoI and XbaI respectively. Each plasmid was purified by phenol:chloroform extraction and ethanol precipitation, resuspended in nuclease-free water. Linearized YFV and WNV plasmids were used for in vitro transcription using the mMessage mMachine SP6 and T7 kits respectively following the manufacturer’s instructions. In vitro transcribed RNAs were purified by phenol:chloroform extraction and ethanol precipitation, diluted to 1 mg/ml, aliquoted and stored at −80C. Vero cells (200,000 cells/well) were transfected with 5 μg of YFV and WNV *in vitro* transcribed RNA via lipofectamine 2000 following the manufacturer’s instructions and incubated at 37°C for 48 hrs. Following incubation, virus titers were quantified by plaque assay as described below. All work with WNV was performed under biosafety level 3 conditions.

#### Deep sequencing analysis

Sequencing data was published previously and can be found on Sequence Read Archive (SRA) under BioProject ID PRJNA589089 [23]. Sequencing reads were processed and aligned as described previously [23]. Minority variants were identified using an in-house variant caller (https://github.com/GhedinLab/ZIKV_Analysis). Coverage and minority variant calls were checked to ensure overlapping regions were identical in their nucleotide composition before merging the fastq files and then realigning the 3 amplicons to the reference file at once. Minority variants were called again on the merged alignment files and the amino acid position was added using the positions indicated on the MR766 NCBI site (KX830961.1). Minority variants were called if coverage at the given nucleotide was at or above 500x and the frequency of the variant was above 1%, present in both forward and reverse reads, and had a quality score above 25.

### Plaque assay and focus-forming assay

Viral titers were determined by plaque assay or focus-forming assay on Vero cells. For plaque assays, virus was subjected to ten-fold serial dilutions in DMEM and added to a monolayer of Vero cells for one hour at 37°C. Following incubation, a 0.8% agarose overlay was added, and cells were incubated for five days at 37°C. Five days post infection, cells were fixed with 4% formalin, the agarose overlay removed, and plaques were visualized by staining with crystal violet (10% crystal violet and 20% ethanol). Viral titers were determined on the highest dilution virus could be counted.

For focus-forming assays (FFA), duplicates of serial 10-fold dilutions of virus in viral growth medium (DMEM containing 2% FBS and 20 mM HEPES) were applied to Vero cells in 96-well plates and incubated for 1 hr at 37°C. Following virus adsorption, the monolayers were overlaid with 1% methylcellulose in minimum essential medium Eagle (MEM). Infected cell foci were detected 42-46 hrs after infection. Following fixation with 2% paraformaldehyde (PFA) for 1 hr at room temperature, plates were incubated with 500 ng/ml of flavivirus cross-reactive mouse MAb E60 [44] for 2 hr at room temperature or overnight at 4°C. After incubation at room temperature for 2 hr with a 1:5,000 dilution of horseradish peroxidase (HRP)-conjugated goat anti-mouse IgG (Sigma), foci were detected by addition of TrueBlue substrate (KPL). Foci were quantified with a CTL Immunospot analyzer.

### Virus growth curves

For multi-step growth analysis, cells were infected at a multiplicity of infection of 0.1 and incubated at 37°C or 28°C with 5% CO_2_ for one hour. Then, inoculum was aspirated and cells were washed with PBS and media was replenished. Samples of infected cell culture supernatant were collected at 4, 24, 48 and 72 hrs post-infection and stored at −80°C for virus titration. Virus quantification was performed by FFA on Vero cells as above.

### Immunoblotting

293T cells (500,000 cells/well in a 6-well plate) were transfected with 1.2 μg of each plasmid as described above. At predetermined time points, cells were harvested, washed twice with PBS, and resuspended in 2x Laemmli buffer containing 10% β-mercaptoethanol. Samples were boiled for 10 mins, centrifuged for 5 min at 12,000 x g to clarify debris, and proteins were separated by SDS-PAGE. Proteins were transferred to a PVDF membrane and membranes were blocked with 5% nonfat milk in Tris-buffered saline (TBS: 50 mM Tris pH 7.5, 150 mM NaCl) containing 0.1% Tween 20 (TBS-T). Membranes were incubated with anti-ZIKV capsid (Genetex; Cat #GTX133317), prM (Genetex; GTX133305), envelope (Genetex; Cat #GTX133314), NS1 (Genetex; Cat #GTX133307), NS3 and NS5 (gifts from Dr. Andres Merits at the University of Tartu, Estonia), and anti-GAPDH (GA1R ThermoFisher; Cat # MA5-15738) antibodies, washed with TBS-T and incubated with a horseradish peroxidase conjugated secondary antibody. After incubation with secondary antibody, membranes were washed extensively, developed with the SuperSignal West Pico chemiluminescent substrate (Pierce), and imaged on the BioRad Chemidoc imager. Images were analyzed and processed using ImageLab (Version 6.0.0).

### Mosquito infections and harvests

*Ae. aegypti* mosquitoes (Poza Rica, Mexico, F30-35) were a gift from Dr. Gregory Ebel at Colorado State University [45]. Mosquitoes were reared and maintained in the NYU School of Medicine ABSL3 facility at 28°C and 70% humidity with a 12:12 hour diurnal light cycle. The day before infection, female mosquitoes were sorted and starved overnight. The day of infection, mosquitoes were exposed to an infectious bloodmeal containing freshly washed sheep blood, 5 mM ATP, and 10^7^ PFU/ml virus. After approximately 30 mins, mosquitoes were cold-anesthetized, and engorged female mosquitoes were sorted into new cups [46] [22]. Engorged mosquitoes were incubated at 28°C with 70% humidity for 14 days and fed ad libitum with 10% sucrose. Following incubation, mosquitoes were cold-anesthetized and the legs and wings removed. Mosquito bodies and legs and wings were put into a 2 ml round-bottom tubes containing 250 ml of PBS and a steel ball (Qiagen). Samples were homogenized using a TissueLyser (Qiagen) with 30 shakes/second for 2 mins, centrifuged at 8,000 x g for 5 mins to remove debris, and viral titers were quantified by plaque assay.

### Mouse infections

4- and 7-day old WT C57BL/6J mouse experiments were completed in the NYU ABSL3 facility performed under the approval of the NYU School of Medicine Institutional Animal Care and Use Committee (IACUC) (Protocol # IA16-01783). 4- and 7-day old male and female C57BL/6J mice were infected subcutaneously with 10^4^ PFU of the WT ZIKV or the ZIKV E^V114M^ variant. Mice were euthanized by decapitation seven dpi, organs harvested, homogenized as described above, and ZIKV titers were quantified by plaque assay.

Husbandry and infections in type I Interferon-receptor knockout (*Ifnar1^-/-^*) mice were performed under the approval of the University of North Carolina at Chapel Hill IACUC (Protocol # 19-185). 5- to 6-week-old male *Ifnar1^-/-^* mice on a C57BL/6 background were used. Mice were inoculated with 10^3^ FFU of ZIKV in a volume of 50 μl by a subcutaneous route in the footpad. Survival and weight loss were monitored for 14 days. Animals that lost ≥30% of their starting weight or that exhibited severe disease signs were euthanized by isoflurane overdose. Weights are reported as percent of weight at the day of infection and group means were censored once one animal in a group died. To measure viral loads, ZIKV-infected mice were euthanized at 4 dpi as above and perfused with 20 ml of PBS. Spleens were harvested and homogenized with zirconia beads (BioSpec) in a MagNA Lyser instrument (Roche Life Science) in 1 ml of PBS. Blood was collected at 2 dpi by submandibular bleed with a 5 mm Goldenrod lancet and by cardiac puncture at 4 dpi. Blood was collected in serum separator tubes (BD) and serum was separated by centrifugation at 8000 rpm for 5 min. Tissues and serum from infected animals were stored at −80°C until titration by focus-forming assay.

### Virus protein accession numbers

GenBank accession numbers used for flavivirus protein alignments: Zika MR766 (KX830960.1), ZIKV Puerto Rican (KX377337), ZIKV Brazilian (KX280026.1), ZIKV French Polynesia (KJ776791.2), ZIKV Cambodian (KU955593.1), DENV1 (BCG29749), DENV2 (BCG29762), DENV3 (AVF19960), DENV4 (BCG29769), YFV (NP_740305), and WNV (MN849176).

### Protein structures

The E glycoprotein structure of ZIKV (PDB: 5JHM), WNV (PDB: 2HG0), YFV (PDB: 6iw4), and the CHIKV E1 glycoprotein structure (PDB: 3n4) were visualized using Protein 3D (www.dnastar.com).

### Statistical analysis

GraphPad Prism (Version 8.0.2) was used to analyze all data and perform statistical analyses. All *in vitro* experiments were completed in triplicate with internal duplicates or triplicates. Mosquito and mouse experiments were completed in two independent infection. p<0.05 is considered significant.

## Acknowledgements

We thank all members of the Stapleford Lab for critical comments on the experiments and manuscript. We thank our collaborators Drs. Julie Pfeiffer, Matthew Evans, Alexander Pletnev, Andres Merits, and Gregory Ebel for essential reagents. This work was supported by a Start-up package from NYU Grossman School of Medicine (K.A.S), the Public Health Service Institutional Research Training Award T32 AI007180 (M.V.R and K.E.E.J), the American Heart Association Postdoctoral Fellowship (19-A0-00-1003686), and R01 AI39512 (H.M.L. and N.C). This work was supported in party by the Division of Intramural Research (DIR) and the NIAID/NIH (E.G.).

